# *Pseudomonas aeruginosa* can diversify after host cell invasion to establish multiple intracellular niches

**DOI:** 10.1101/2022.10.07.511388

**Authors:** Naren G. Kumar, Vincent Nieto, Abby R. Kroken, Eric Jedel, Melinda R. Grosser, Mary E. Hallsten, Matteo M. E. Mettrucio, Timothy L. Yahr, David J. Evans, Suzanne M. J. Fleiszig

## Abstract

Within epithelial cells, *Pseudomonas aeruginosa* depends on its type three secretion system (T3SS) to escape vacuoles and replicate rapidly in the cytosol. Previously, it was assumed that intracellular subpopulations remaining T3SS-negative (and therefore in vacuoles) were destined for degradation in lysosomes, supported by data showing vacuole acidification. Here, we report in both corneal and bronchial human epithelial cells that vacuole associated-bacteria can persist, sometimes in the same cells as cytosolic bacteria. Using a combination of phase-contrast, confocal, and correlative light and electron microscopy, we also found they can demonstrate biofilm-associated markers: *cdrA* and cyclic-di-GMP (c-di-GMP). Vacuolar-associated bacteria, but not cytosolic counterparts, tolerated the cell-permeable antibiotic ofloxacin. Surprisingly, use of mutants showed that both persistence in vacuoles and ofloxacin tolerance were independent of the biofilm-associated protein CdrA or exopolysaccharides (Psl, Pel, alginate). A T3SS mutant (Δ*exsA*) unable to escape vacuoles phenocopied vacuolar-associated sub-populations in wild-type PAO1-infected cells, results revealing that epithelial cell death depended upon bacterial viability. Intra-vital confocal imaging of infected mouse corneas confirmed that *P. aeruginosa* formed similar intracellular sub-populations within epithelial cells *in vivo*. Together, these results show that *P. aeruginosa* differs from other pathogens by diversifying intracellularly into vacuolar and cytosolic sub-populations that both contribute to pathogenesis. Their different gene expression and behavior (e.g., rapid replication versus slow replication/persistence) suggest cooperation favoring both short- and long-term interests and another potential pathway to treatment failure. How this intracellular diversification relates to previously described “acute versus chronic” virulence gene-expression phenotypes of *P. aeruginosa* remains to be determined.

**Importance:** *Pseudomonas aeruginosa* can cause sight- and life-threatening opportunistic infections, and its evolving antibiotic resistance is a growing concern. Most *P. aeruginosa* strains can invade host cells, presenting a challenge to therapies that do not penetrate host cell membranes. Previously, we showed that the *P. aeruginosa* type III secretion system (T3SS) plays a pivotal role in survival within epithelial cells, allowing escape from vacuoles, rapid replication in the cytoplasm, and suppression of host cell death. Here, we report the discovery of a novel T3SS-negative sub-population of intracellular *P. aeruginosa* within epithelial cells that persist in vacuoles rather than the cytoplasm, and that tolerate a cell-permeable antibiotic (ofloxacin) that is able to kill cytosolic bacteria. Classical biofilm-associated markers, although demonstrated by this sub-population, are not required for vacuolar persistence or antibiotic tolerance. These findings advance our understanding of how *P. aeruginosa* hijacks host cells, showing it diversifies into multiple populations with T3SS-negative members enabling persistence whilst rapid replication is accomplished by more vulnerable T3SS-positive siblings. Intracellular *P. aeruginosa* persisting and tolerating antibiotics independently of the T3SS or biofilm-associated factors could present additional challenges to development of more effective therapeutics.

## INTRODUCTION

*Pseudomonas aeruginosa* is a significant cause of morbidity and mortality including burn-wound infections, septicemia, catheter-associated infections, corneal infections, community-acquired and nosocomial pneumonia, and chronic lung disease in persons with cystic fibrosis (CF). *P. aeruginosa* infections are notoriously difficult to treat due to a combination of inherent antimicrobial resistance and the versatility and adaptability of this opportunistic pathogen (1–5). While often considered an extracellular pathogen, several decades of work across many research labs have established that clinical isolates of *P. aeruginosa* can internalize and survive in host cells, including in animal models of infection (6–14). Despite this, surprisingly little is known about the intracellular lifestyle of *P. aeruginosa*, and many questions remain about how it contributes to disease pathogenesis and recalcitrance to therapeutics.

Previously, we showed that *P. aeruginosa* survival inside cells is modulated by the type three secretion system (T3SS) and several of its effectors (ExoS, ExoT and ExoY), with ExoS playing a prominent role (11, 15, 16). While the RhoGAP activity of ExoS (and ExoT) can counter bacterial internalization by host cells via antiphagocytic activity (15, 17–20), bistability results in a lack of T3SS expression in many extracellular bacteria, effectively tempering antiphagocytic activity and allowing some population members to invade cells (11, 16, 21). Once inside the cell, efficient T3SS triggering allows expression of the T3SS, which in an ExoS-dependent manner inhibits vacuole acidification, enables escape from the endocytic vacuole, inhibits autophagy, and allows rapid replication of bacteria in the host cytosol (10, 17, 22, 23). Differing from other intracellular bacteria that use host cytoskeletal components for intracellular motility (24), *P. aeruginosa* then utilizes its own pili/twitching motility to disseminate throughout the host cell (25). Meanwhile, ExoS further supports intracellular survival by delaying lytic host cell death effectively preserving the intracellular niche (16), and it drives formation of plasma membrane blebs to which some bacteria traffic and replicate within (9, 19). These blebs can subsequently disconnect from cells, becoming vesicles with intact membranes that can carry enclosed live/swimming bacteria to distant sites (9). Mutations affecting T3SS needle assembly, T3SS toxin secretion, or expression of the entire T3SS (Δ*exsA*), restrict intracellular bacteria to endocytic vacuoles (10, 11, 15), which we previously presumed were destined for degradation within lysosomes (10).

In a previous study, we showed that mutants defective in twitching motility, and therefore unable to disseminate intracellularly, formed non-motile bacterial aggregates inside host cells surrounded by electron-lucid halos on electron micrographs (25). This led us to explore if *P. aeruginosa* was able to produce biofilm-associated factors inside infected host cells.

Biofilm formation has long been recognized as a strategy that bacteria use to survive under adverse environmental conditions including during infections (26, 27). In human infections, *P. aeruginosa* biofilm formation can be associated with chronic persistence in the host, through a combination of phenotypic and genotypic adaptations that allow acquisition of essential nutrients and confer resistance to antimicrobial therapies and host defenses (28–31). Investigations of *P. aeruginosa* biofilm architecture and composition have revealed the presence multiple bacterial exopolysaccharides in the biofilm matrix including Psl, Pel and alginate, the latter most prominent in *P. aeruginosa* biofilms in CF (32, 33). CdrA is a bacterial protein with production regulated by the bacterial second messenger c-di-GMP that reinforces and protects the biofilm matrix (34, 35). Extracellular DNA (eDNA) is another major component of the biofilm matrix that interacts with polysaccharide components and can modulate bacterial dispersal (36–38).

To test the hypothesis that intracellular *P. aeruginosa* could demonstrate biofilm-associated markers we used gene expression reporters for *cdrA* encoding the biofilm-matrix protein and c-di-GMP (which triggers biofilm formation). Results for both corneal and bronchial epithelial cells showed distinct intracellular populations when cells were infected with wild-type *P. aeruginosa*, with *cdrA*-expressing bacteria localized to vacuoles and T3SS-expressing bacteria in the cytosol, often in the same cell. The data also showed potentially important functional differences between the two populations, with cytosolic *P. aeruginosa* sensitive to the cell permeable antibiotic ofloxacin, and vacuolar populations demonstrating ofloxacin tolerance even at high concentrations. Surprising, neither persistence in vacuoles nor ofloxacin tolerance required *cdrA* or any of the three exopolysaccharides (*alg, psl, pel*) known to be associated with biofilm formation. Raising potentially interesting implications for the impact of antibiotics on infection pathogenesis, the results also showed that the vacuolar-located antibiotic survivors were able to trigger cell death.

The discovery that *P. aeruginosa* can establish niches in multiple locations within the same epithelial cells (cytoplasm, membrane blebs, vacuoles) differs from other bacterial pathogens which traffic to either vacuoles or the cytoplasm in a linear fashion. The finding that the bacteria in these alternate locations express different survival/virulence determinants and vary in antibiotic susceptibility predicts a “covering of the bases” for the overall population. Interestingly, this pattern of gene expression occurring simultaneously in infected cells is reminiscent of the “acute” versus “chronic” infection expression phenotypes, which are generally thought to represent entirely different infection types. How intracellular diversification relates to those previously described infection phenotypes, and the relevance of these results to antibiotic treatment failure *in vivo*, will require further investigation.

## RESULTS

### Bacteria expressing biofilm-associated factors *cdrA*, c-di-GMP localize to vacuoles in human epi thelial cells

To determine if biofilm-associated factors were expressed by intracellular *P. aeruginosa*, human corneal and bronchial epithelial cells were imaged 6 h after inoculation with *P. aeruginosa* that report expression of *cdrA* (pMG078) or c-di-GMP (pFY4535) (39, 40) (Table 1). Results were compared to bacteria that instead report T3SS expression (10, 11). Following a 3 h infection period to allow bacteria to invade cells, extracellular bacteria were eliminated using the non-cell permeable antibiotic amikacin, thereby allowing intracellular bacteria to be selectively visualized (8, 10, 11).

**Table 1.** *P. aeruginosa* strains, mutants and plasmids used in this study

| Name |  | Description | Source |
| --- | --- | --- | --- |
| **Strain or Mutant** |  |  |  |
| PAO1F |  | Wild-type | Dr. Alain Filloux |
| PAO1FD$exsA$ | | $exsA$ mutant | Dr. Alain Filloux |
| PAO1P |  | Wild-type | Dr. Matthew Parsek (34) |
| PAO1P$\Delta cdrA$ | | $cdrA$ mutant | Dr. Matthew Parsek (34) |
| PAO1P$\Delta EPS$ | | $psl$, $pel$, $algD$ mutant | Dr. Matthew Parsek (32) |
| PAO1P$\Delta EPS\Delta cdrA$ | | $psl$, $pel$, $algD$, $cdrA$ mutant | Dr. Matthew Parsek (32) |
| mPAO1 |  | Wild-type PAO1 | *Pseudomonas* transposon library (72) |
| **Plasmid** |  |  |  |
| pMG078 | *pBBR1 oriV* | $cdrA$-GFP | This study |
| pFY4535 |  | c-di-GMP<br>Turbo-RFP reporter | Dr. Fitnat Yildiz (39. 40) |
| pJNE05 | *Plasmid:*<br>*pJN105*, *ori:*<br>*pBBR1oriV* | T3SS-GFP expression<br>vector<br>(pJNE05::$exoS$-GFP) | Dr. Timothy Yahr (73) |
| pBAD-GFP<br>(pGFP<sub>arabinose</sub>) | *Plasmid:*<br>*pJN105*, *ori:*<br>*pBBR1oriV* | Arabinose-inducible GFP<br>(pTJ1::$araGFP$) | This study |
| pMG055 | *Plasmid:*<br>pTJ1, *ori:*<br>*oriT* | Constitutive blue<br>fluorescent protein<br>(EBFP2) | This study |

Aligning with our previously published data (9–11, 16), T3SS-expressing intracellular bacteria localized to the cytosol in both epithelial cell types, wherein they replicated rapidly and disseminated throughout the cytoplasm (Fig. 1). Bacteria expressing the biofilm-associated factors *cdrA-* or c-di-GMP-were also detected among the intracellular population. Rather than being in the cytosol, this sub-population appeared to be contained in vacuolar-like niches that remained stable over time (Fig. 1A). To confirm that *cdrA*-GFP reporting bacteria (pseudo-colored red) were within vacuoles as opposed to being in aggregates within the cytosol, real-time phase-contrast microscopy was used to visualize the cellular compartment in which they were contained. Fig. 1B shows *cdrA*-GFP reporting PAO1F (11, 16, 21) located in vacuoles in corneal epithelial cells and Fig. 1C shows the same PAO1F phenotype in bronchial epithelial cells. A different wild-type parental strain PAO1P (35, 41) (see Table 1) also showed the same *cdrA*-GFP reporting vacuolar-located phenotype within corneal epithelial cells (data not shown). Thus, for two PAO1 parental [wild-types, some of the sub-population expressing *cdrA* were found restricted within membrane-bound circular phase-clear regions implicating vacuoles and distinguishing them from possible intracellular cytosolic aggregates.

**Figure 1.**
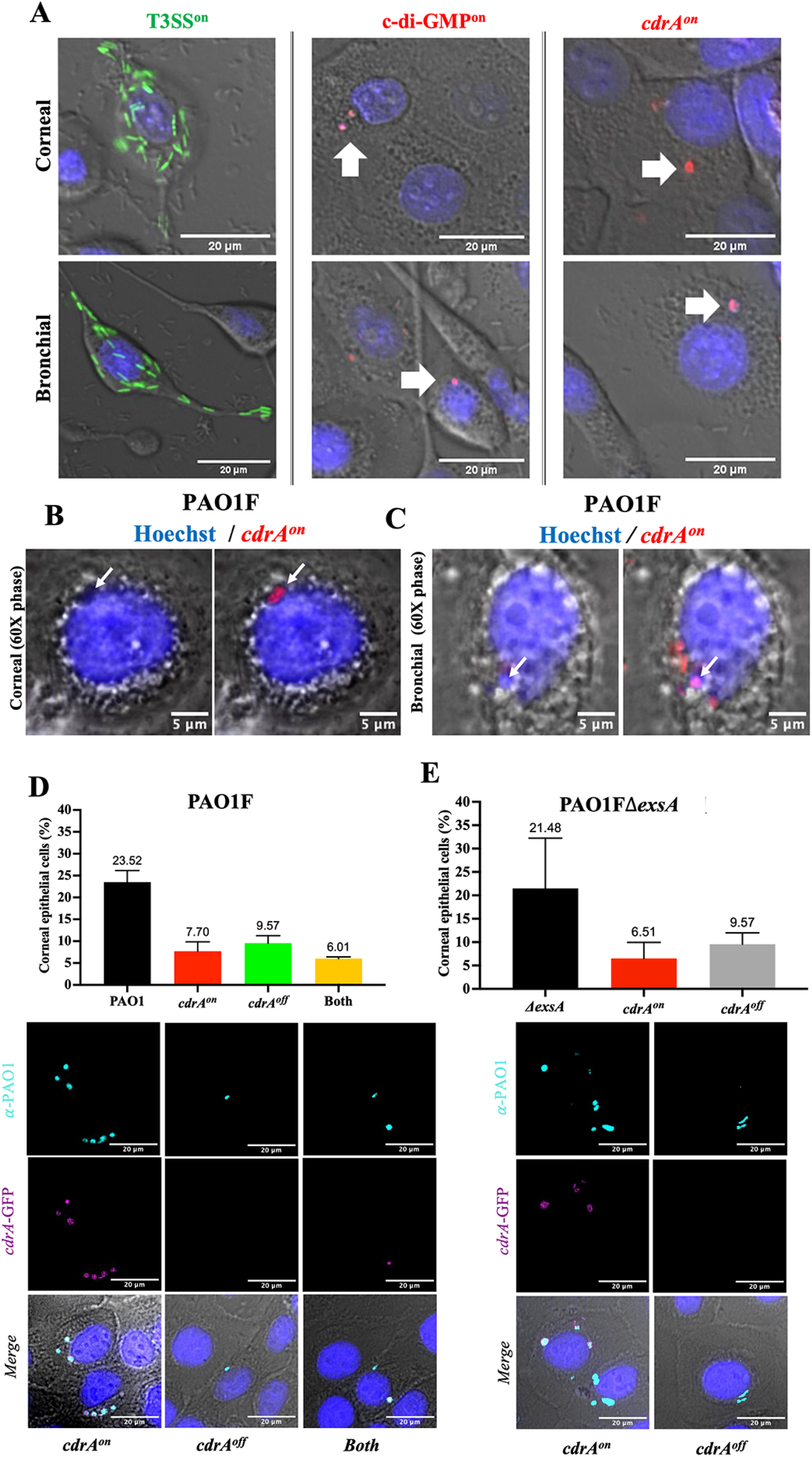
Intracellular wild-type *P. aeruginosa* form distinct phenotypes within human epithelial cells. A) Intracellular expression of genes encoding the T3SS (green), chronic switch regulator c-di-GMP (red), or biofilm matrix protein CdrA (*cdrA*-GFP, pseudo-colored red) in human corneal or bronchial epithelial cells at 6 h post-infection with *P. aeruginosa* PAO1F. Extracellular bacteria were killed at 3 h post-infection with amikacin (200 μg/mL). T3SS-expressing bacteria occupied the cytosol as expected. In contrast, bacteria reporting c-di-GMP or *cdrA*-expression formed distinct localized vacuole-like niches within the same or different cells. B) and C) 60 X oil-immersion phase-contrast microscopy shows *cdrA*-GFP reporting bacteria (pseudo-colored red puncta) of wild-type PAO1F localized to vacuoles in corneal epithelial cells (B) or bronchial epithelial cells (C). D) Percentage of corneal epithelial cells containing any intracellular *P. aeruginosa* (PAO1F, black bar) and distinct *cdrA*-expressing phenotypes. After labeling all bacteria with anti-*Pseudomonas* antibody, *cdrA*-GFP reporter expression allowed sub-division of invaded cells into those only containing *cdrA*^on^ bacteria (red bar), only *cdrA*^off^ (green bar), and those with both *cdrA*^on^ and *cdrA*^off^ (yellow bar). E) Percentage of corneal epithelial cells containing intracellular PAO1FΔ*exsA* (black bar), a T3SS mutant that only localizes to vacuoles (9). Cells containing *exsA* mutants showed a similar vacuolar distribution to wild-type with some only containing *cdrA*^on^ (red bar) and others containing *cdrA*^off^ vacuolar bacteria (grey bar). In (D) and (E), there was no significant difference between sub-groups in distribution of vacuolar phenotypes (One-way ANOVA with Dunnett’s multiple comparisons test). Data are shown as the mean +/-SD of three biological replicates. Images below quantitative data in (D) and (E) show examples of corneal epithelial cells for each of the categories quantified (bacteria pseudo-colored as indicated: *cdrA*-GFP - magenta, anti-*Pseudomonas* antibody - cyan).

Having shown distinct intracellular phenotypes of *P. aeruginosa*, the percentage of corneal epithelial cells containing *cdrA*-expressing intracellular bacteria was then quantified. Cells were infected with PAO1F expressing the *cdrA*-GFP reporter (pMG078) and total intracellular bacteria detected with an anti-*Pseudomonas* antibody at 6 h post-infection (Fig, 1D). The percentage of cells containing intracellular bacteria that expressed only *cdrA (cdrA^on^*) was quantified, along with the percentage of cells containing intracellular bacteria not expressing *cdrA* (*cdrA*^off^), or those containing both. Of 520 cells analyzed, 7.7 % contained only *cdrA*-expressing bacteria (red bar), and 9.6 % contained *cdrA*^off^ bacteria (green bar), while 6 % contained both phenotypes (yellow bar) (Fig. 1D). Images below quantitative data show examples of each of these different categories.

To determine if a lack of vacuolar escape into the cytosol necessarily leads to *cdrA* expression, we performed the same analysis using a mutant lacking ExsA, the transcriptional activator of the T3SS (PAO1FΔ*exsA*) (9). This mutant does not express the T3SS and remains in vacuoles because the T3SS is required for vacuolar escape and survival in the cytosol (10). Results (Fig. 1E) confirmed that at this time point (6 h post-infection), *exsA*-mutants were only present in vacuoles and that the percentage of cells containing internalized bacteria was similar to wild-type parent PAO1F (see Fig. 1D). Importantly, the distribution of *cdrA*^on^ versus *cdrA*^off^ bacteria found in vacuoles for *exsA*-mutant infected cells (Fig. 1E) was similar to wild-type-infected cells (see Fig 1D). Images of *cdrA*^on^ and *cdrA*^off^ categories for *exsA*-mutant infected cells are shown (Fig. 1E).

Thus, both wild-type bacteria and T3SS (*exsA*) mutants can express *cdrA* when in vacuoles, and this occurs in a similar percentage of the infected host cell population under these experimental conditions.

### Vacuolar localization does not require *cdrA*

C-di-GMP is a trigger for biofilm formation while CdrA is the only known biofilm matrix protein made by *P. aeruginosa*. Since *cdrA* was expressed only in bacteria localized to vacuoles, we next asked if vacuolar localization depends on CdrA, and potentially also on biofilm formation. Thus, we compared a *cdrA* mutant (PAO1PΔ*cdrA*) to wild-type (PAO1P) for the ability to persist in vacuoles. First, we confirmed that the *cdrA* mutant did not differ from wild-type in its ability to invade corneal epithelial cells (Fig. 2A) and did not have a different impact on cell viability over time (data not shown). Next, we compared the percentage of corneal epithelial cells that contained at least one *cdrA-*GFP expressing bacterium after infection with wild-type (PAO1P) or *cdrA* mutant (PAO1PΔ*cdrA*) at 6 h post-infection. Results showed that the *cdrA* mutant and wild-type trafficked to vacuoles in a similar percentage of cells over the 6 h period (Fig. 2B). Phase-contrast microscopy also showed that mutants unable to express *cdrA* (Δ*cdrA*) trafficked to vacuoles at 6 h post-infection (Fig. 2C). This further shows that vacuolar persistence at 6 h post-infection did not require CdrA (Fig. 2B and C).

**Figure 2.**
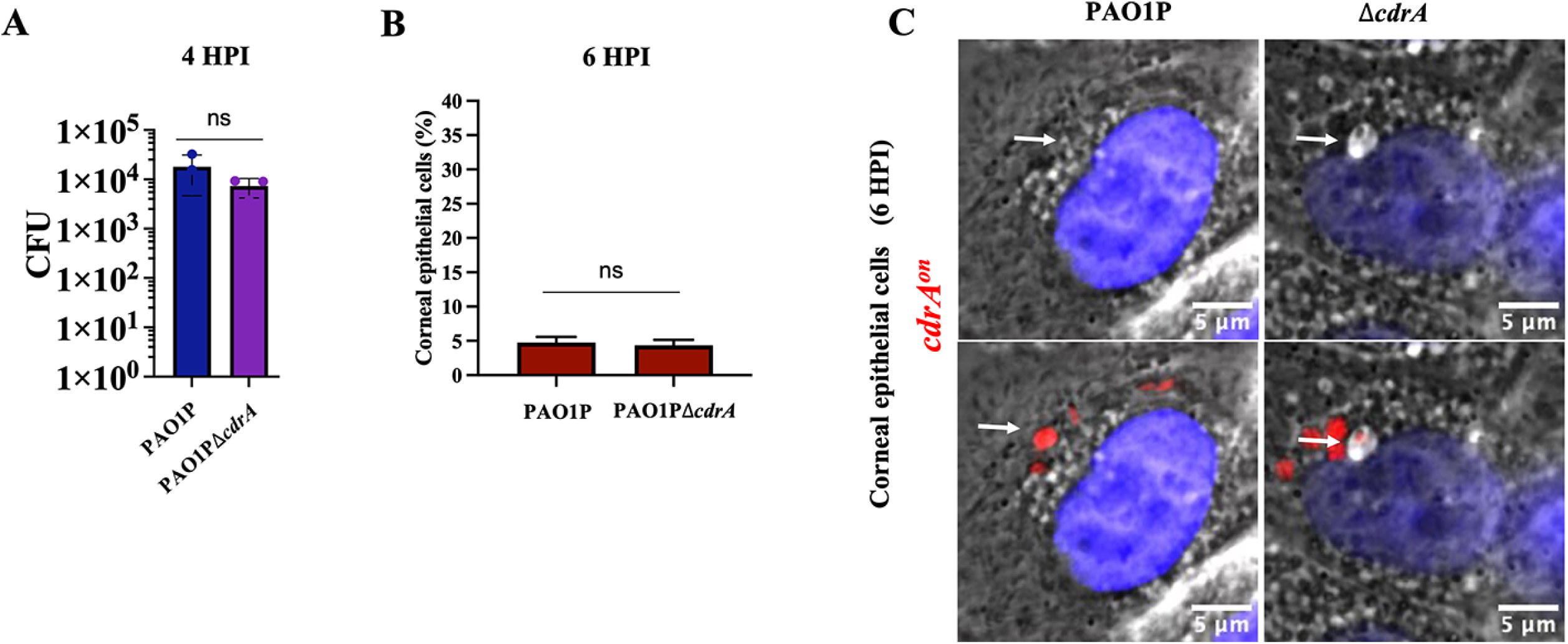
CdrA is not required for host cell invasion and vacuolar localization. A) Invasion of corneal epithelial cells quantified by quantifying colony forming units (CFU) at 4 h post-infection. Corneal cells were infected for 3 h with PAO1P and PAO1PΔ*cdrA*. Extracellular bacteria were killed 3 h post-infection with amikacin (200 μg/mL) and cells lysed to recover intracellular bacteria. B) Percentage of human corneal epithelial cells containing at least one *cdrA*^on^ bacterial cell at 6 h post-infection. C) 60 X oil-immersion phase-contrast microscopy shows *cdrA*-expressing bacteria (red puncta) of wild-type PAO1P and its mutant PAO1PΔ*cdrA* localized to vacuoles in corneal or bronchial cells. White arrows point to bacteria in vacuoles. The same image is shown with and without fluorescence. ns = Not Significant (Student’s t-Test).

### Among intracellular bacteria, the *cdrA-*expressing population is more tolerant to ofloxacin than the T3SS-expressing population

Biofilm-associated bacteria tend to be more antibiotic tolerant. Thus, we next explored if the intracellular *cdrA*-expressing population could better resist an antibiotic than their cytosolic T3SS-expressing cell-mates. This was done using ofloxacin, an antibiotic differing from amikacin (in part) by being able to penetrate host cell membranes. Wild-type bacteria were allowed to internalize for 3 h, before exposing epithelial cells to either amikacin alone (non-cell permeable, kills only extracellular bacteria) or amikacin plus ofloxacin (to additionally target intracellular bacteria). Ofloxacin concentrations were chosen according to the MIC for the PAO1 strain variants used (see methods). For PAO1F, the MIC for ofloxacin was determined to be 0.25 μg/mL, and for that reason concentrations used ranged from 0.25 - 4 μg/mL (1X-16X MIC). Fig. 3 shows time-lapse imaging of intracellular *P. aeruginosa* gene expression using T3SS-GFP and *cdrA*-GFP reporters, and the percentage of epithelial cells containing at least one GFP-positive bacterial cell at 6 and 12 h post-infection.

**Figure 3.**
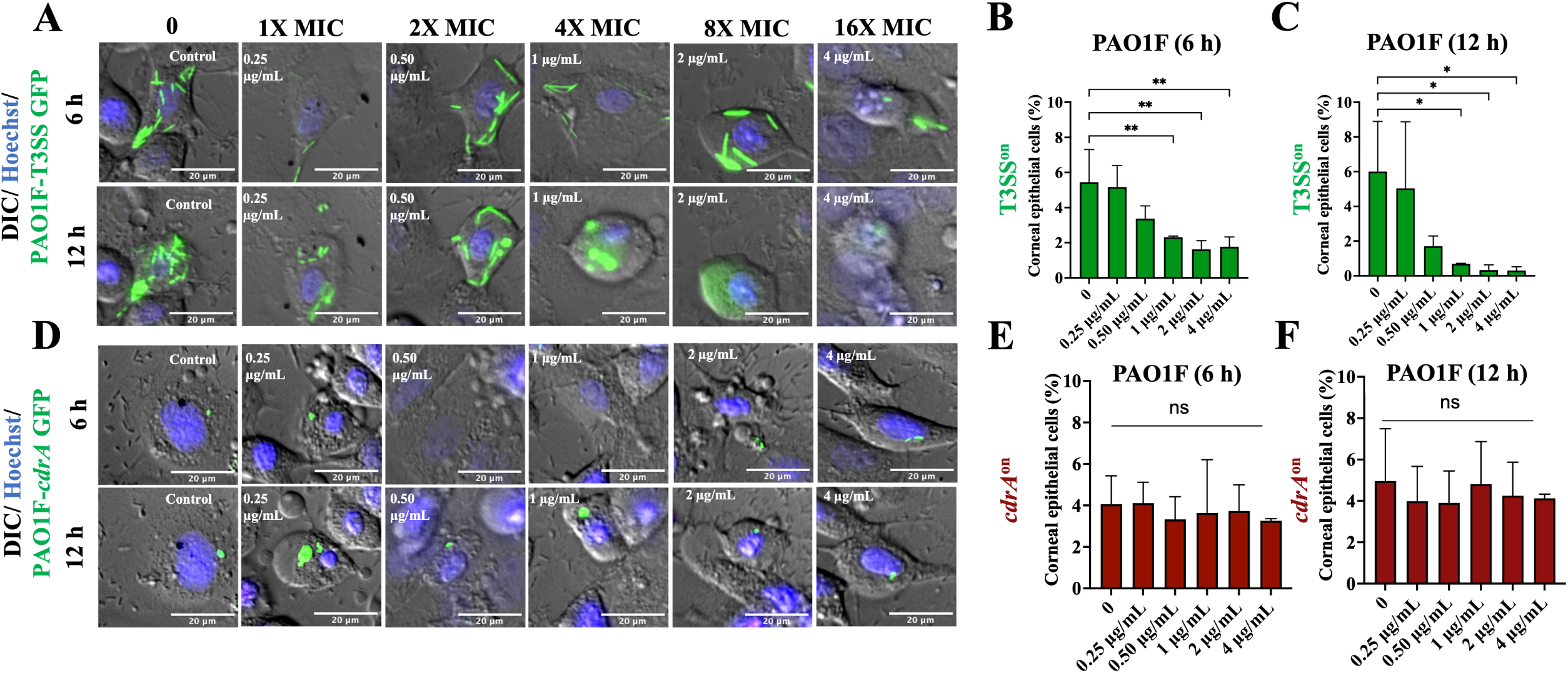
Vacuolar *cdrA*^on^ populations show resistance to the cell permeable antibiotic ofloxacin compared to cytosolic T3SS^on^ bacteria. A) Images of human corneal epithelial cells containing cytosolic T3SS-GFP-expressing PAO1F (T3SS^on^) in the presence of ofloxacin (0.25 - 4 μg/mL = 1 - 16X MIC) at 6 h and 12 h post-infection versus control. Extracellular bacteria were killed after 3 h with the non-cell permeable amikacin (200 μg/mL) (control) or amikacin (200 μg/mL) and ofloxacin were added at 3 h to also target intracellular bacteria. B-C) Percentage of corneal epithelial cells containing at least one T3SS^on^ bacterial cell at 6 h (B) and 12 h (C) post-infection under the above conditions. Increasing concentrations of ofloxacin eliminated most cytosolic bacteria (* p < 0.05, ** p < 0.01, ns = Not Significant, One-way ANOVA with Dunnett’s multiple comparisons test). D) Images of *cdrA*-GFP-expressing PAO1F vacuolar bacteria (*cdrA*^on^) in the presence of ofloxacin as above. Discrete foci of bacteria (green) persisted at 16X the MIC of ofloxacin and at 12 h post-infection. E-F) Percentage of human corneal epithelial cells containing at least one *cdrA*^on^ bacterial cell at 6 h (E) and 12 h (F) post-infection under the above conditions. Increasing concentrations of ofloxacin failed to eliminate the *cdrA*^on^ vacuolar population. No significant difference between groups (One-way ANOVA with Dunnett’s multiple comparisons test). Data shown as mean +/- SD of three biological replicates.

Cytosolic T3SS^on^ populations showed significant susceptibility to ofloxacin at 1 μg/mL (4X MIC) at both 6 h and 12 h, with fewer infected cells detected over time (Fig. 3A, B, C and Supplemental Video S1). In contrast, intracellular *cdrA*^on^ bacteria continued to report with ofloxacin treatment concentrations all the way up to 4 μg/mL (16X MIC) at both 6 and 12 h post-infection with ∼ 4-6 % of epithelial cells containing *cdrA*^on^ bacteria (Fig. 3D, E, F and Supplemental Video S2). These results showed that vacuolar *cdrA*-expressing bacteria were better able to resist killing by ofloxacin than T3SS-reporting cytosolic bacteria in the same infected host cell population.

### Correlative light and electron microscopy confirmed viability and vacuolar localization of ofloxacin tolerant *P. aeruginosa*

Since the fluorescent signal from intracellular bacteria may persist long after bacterial cell death (42) and *cdrA* expression is induced upon cell contact, correlative light and electron microscopy (CLEM) was used to confirm that *cdrA*-GFP reporting bacteria surviving above MIC concentrations of ofloxacin were both intracellular and localized to vacuoles. CLEM is essentially a combination of wide-field immunofluorescence and electron microscopy (EM) using the same section to localize fluorescence with morphological details (e.g. of a cell) (43, 44). Ofloxacin was used at 6 h post-infection (with PAO1F) at a concentration that can clear the cytosolic population (1 μg/mL, see Fig. 3A). Fig. 4 shows three individual cells containing *cdrA*-GFP expressing bacteria in vacuoles, all localized to the perinuclear region where late endosomes are expected to reside, each containing ∼1 - 4 bacterial cells. Since vacuolar localized bacteria have intact cell walls after antibiotic treatment, it suggests that vacuolar-localized populations are viable. This outcome supports the conclusion from our wide-field fluorescence and DIC/phase-contrast microscopy experiments (Fig. 1A, B; Fig. 2C; Fig. 3D), that ofloxacin tolerant *cdrA*-expressing sub-populations can be intracellular *P. aeruginosa* and that they localize to vacuoles.

**Figure 4.**
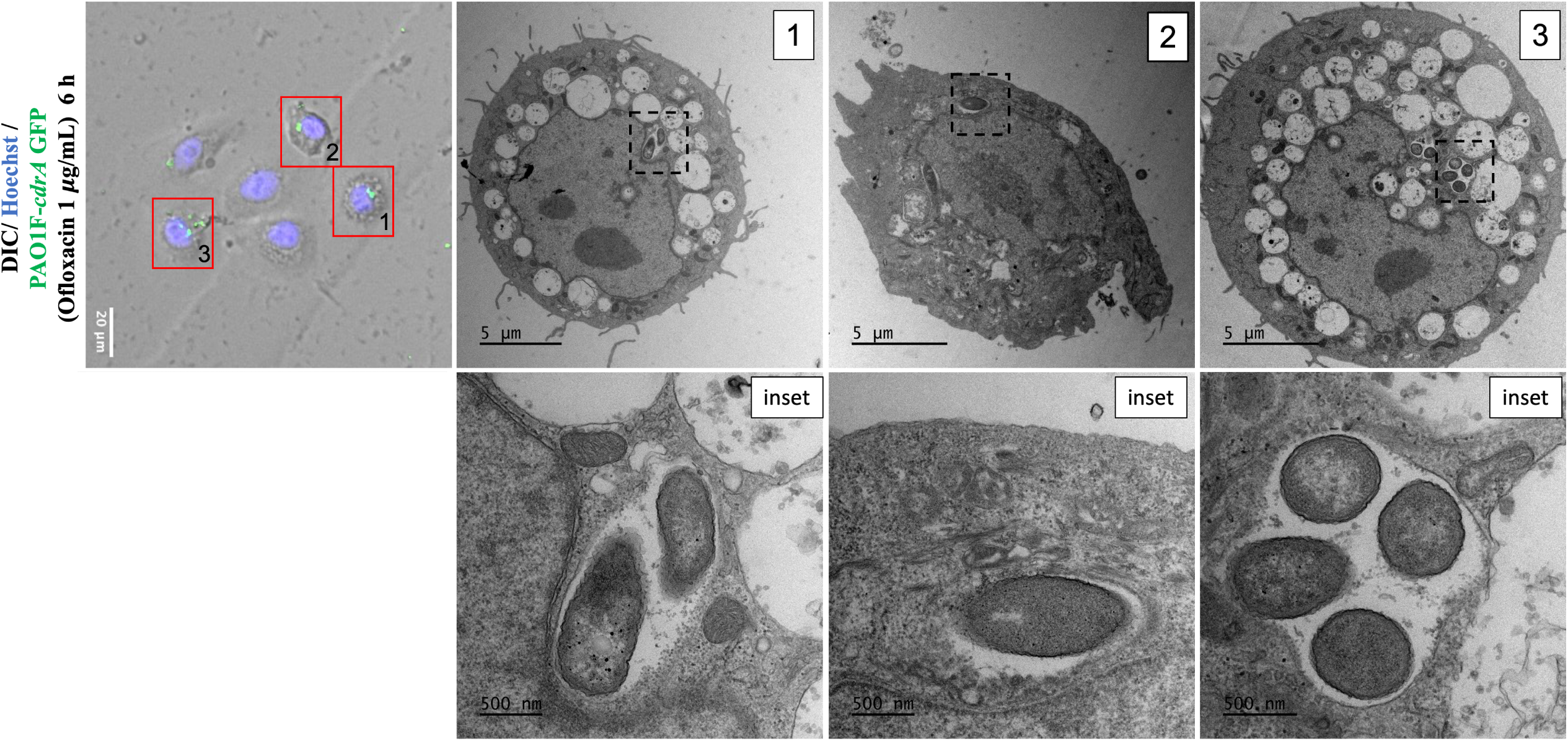
Correlative light and electron microscopy of *cdrA*^on^ bacteria in vacuoles. Fluorescence and electron micrographs of human corneal epithelial cells at 6 h after inoculation with PAO1F expressing *cdrA*-GFP. Extracellular bacteria were killed 3 h post-infection with amikacin (200 μg/mL) and amikacin (200 μg/mL) and ofloxacin (1 μg/mL) then added for a further 3 h. Cells were fixed at 6 h and fluorescence microscopy performed before preparing cells for electron microscopy (EM). Red boxes show corneal epithelial cells with corresponding EM micrographs. Black dotted boxes indicate regions from which respective insets were taken to show bacteria within vacuoles with evidence of vacuolar membranes.

### Detectable *cdrA* expression does not correlate with ofloxacin resistance

While the experiments above show that *crdA*-expressing intracellular bacteria localized to vacuoles, not all vacuolar-located bacteria expressed *cdrA*. Heterogeneity among vacuolar bacteria is not surprising, as they likely transition through various phenotypes after internalization/replication and as maturation of the vacuole alters their environment. Here, we sought to explore ofloxacin susceptibility among all vacuolar bacteria in a wild-type infection, not just those expressing *cdrA*. To specifically label (all intracellular bacteria) and study the effect of ofloxacin on the total population of intracellular bacteria, ofloxacin was used to kill the susceptible population in cells infected with wild-type bacteria expressing an arabinose-inducible reporter (pGFP_arabinose;_ see Methods & Supplemental Fig. S1) where only the survivors express GFP. Epithelial cells containing at least one ofloxacin survivor were then quantified at 6 and 12 h (Fig. 5). As expected, when ofloxacin was not used or used only at sub-lethal concentrations (< MIC 0.50 µg/mL), epithelial cells infected with wild-type bacteria were found to contain both cytosolic and vacuolar-localized bacteria (Fig. 5A). When ofloxacin was used 2X above the MIC (0.5 μg/mL) or higher, there was a gradual reduction in the number of cytosolic bacteria while only vacuolar populations continued to persist (Fig. 5A). The percentage of cells containing vacuoles with ofloxacin tolerant bacteria was similar at 6 h (Fig. 5B) and 12 h (Fig. 5C), showing stability during the intervening time. Comparison to percentage of cells containing any ofloxacin survivors in vacuoles to those containing only *cdrA*^on^ survivors (∼10 %, Fig. 5B and C versus ∼ 4 %, Fig. 3E and F) suggested additional ofloxacin tolerant bacteria (in vacuoles) besides those expressing *cdrA* (Fig. 5A versus 3D). These results suggested that while vacuolar bacteria can express *cdrA* and can also tolerate high doses of ofloxacin, while expression of *cdrA* is dispensable for vacuolar localization and tolerating high concentrations of the antibiotic (Fig 2C). Unfortunately, experiments to test this more directly could not be done using bacteria simultaneously expressing both reporters (i.e., arabinose-inducible GFP and *cdrA*) because expression of dual fluorescent reporters in *P. aeruginosa* impairs bacterial fitness and affects the invasion of host cells. Instead, we used isogenic *cdrA* mutants as discussed below.

**Figure 5.**
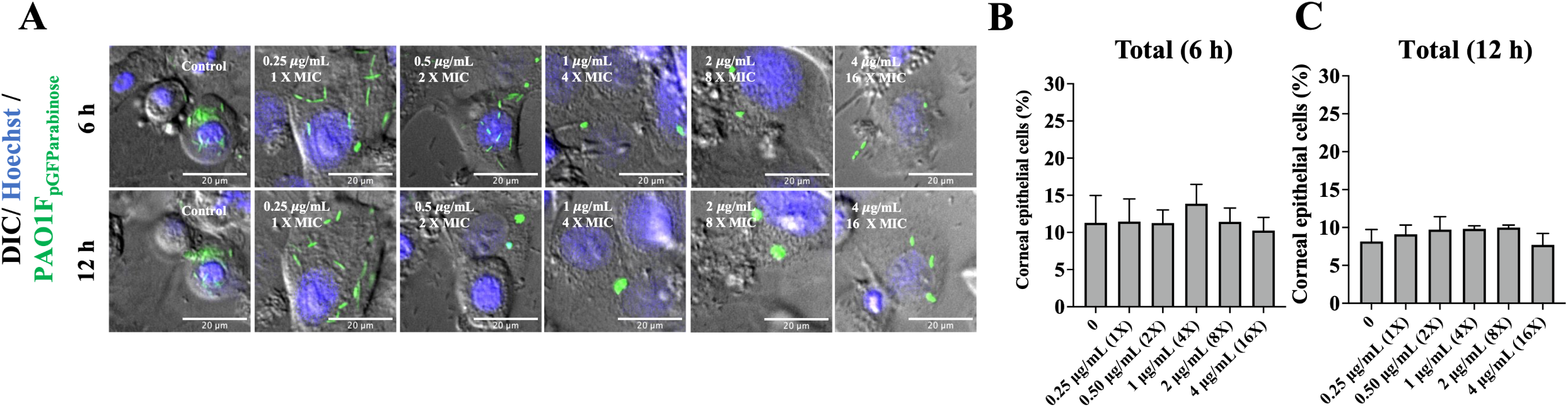
The total population of ofloxacin-tolerant wild-type *P. aeruginosa* localizes to vacuoles. A) Images of human corneal epithelial cells containing PAO1F expressing pGFP_arabinose_ to show all intracellular bacteria in the presence of ofloxacin (0.25 - 4 μg/mL = 1 - 16X MIC) at 6 and 12 h post-infection versus control. After 3 h extracellular bacteria were killed with amikacin (200 μg/mL) (control) or amikacin (200 μg/mL) and ofloxacin were added to also target intracellular bacteria. Cytosolic PAO1F were cleared by ofloxacin at 1 μg/mL at 6 h and by 0.5 μg/mL at 12 h. Vacuolar bacteria persisted in the epithelial cells at 12 h post-infection at all ofloxacin concentrations. B-C) Percentage of corneal epithelial cells containing at least one PAO1 with pGFP_arabinose_ bacterial cell at 6 h (B) and 12 h (C) post-infection under the above conditions. Despite ofloxacin clearance of cytosolic bacteria, the percentage of cells containing intracellular bacteria did not significantly differ between control and ofloxacin treated groups (One-way ANOVA with Dunnett’s multiple comparisons test). Data are shown as the mean +/- SD of three biological replicates.

### CdrA and biofilm-associated EPS is dispensable for ofloxacin resistance of vacuolar *P. aeruginosa*

Having shown expression of the biofilm-associated gene *cdrA* in a subset of vacuolar bacteria, along with vacuolar bacteria as being more tolerant to ofloxacin than cytosolic bacteria, and that ofloxacin tolerant bacteria could express *cdrA*, we next explored if CdrA was required for ofloxacin resistance of vacuolar bacteria. To more broadly study the contribution of biofilm formation, we also considered biofilm-associated exopolysaccharides (EPS). Thus, mutants in CdrA (Δ*cdrA*), exopolysaccharides (Δ*EPS*; *psl, pel, and* alginate) or both (Δ*cdrA*/Δ*EPS)* were tested for intracellular ofloxacin resistance using time-lapse imaging as described above. Since the *in vitro* MIC of PAO1P (the wild-type parent of these mutants) was 4 μg/mL versus 0.25 μg/mL for PAO1F (used above) the ofloxacin concentration was adjusted accordingly from 0 - 64 μg/mL (0 – 16X MIC). Fig. 6 A-D shows the percentage of corneal epithelial cells containing at least one GFP-expressing bacterial cell at 12 h post-infection. Use of p*GFP*_arabinose_ detected all viable intracellular *P. aeruginosa*, while *cdrA*-GFP specifically reported biofilm-associated sub-populations. Surprisingly, there was no significant difference between wild-type and *cdrA* mutants in the number of cells still containing viable intracellular bacteria after ofloxacin treatment at all concentrations up to 64 μg/mL (16X MIC) (Fig. 6A and B) with each strain entering ∼15-20 % of the corneal epithelial cells (Fig. 6A and B, grey bars) and *cdrA*-GFP expressing vacuolar sub-populations in ∼5-10 % of those cells (Fig. 6A and B, red bars). This was shown using arabinose-inducible GFP to detect all intracellular ofloxacin survivors, and the *cdrA*-reporter to detect only the subset reporting *cdrA* promotor activity (which still occurs in *cdrA* mutants) (Fig. 6B). Thus, *cdrA* is not required for the ofloxacin resistance exhibited by vacuolar *P. aeruginosa* in addition to not being required for vacuolar localization.

**Figure 6.**
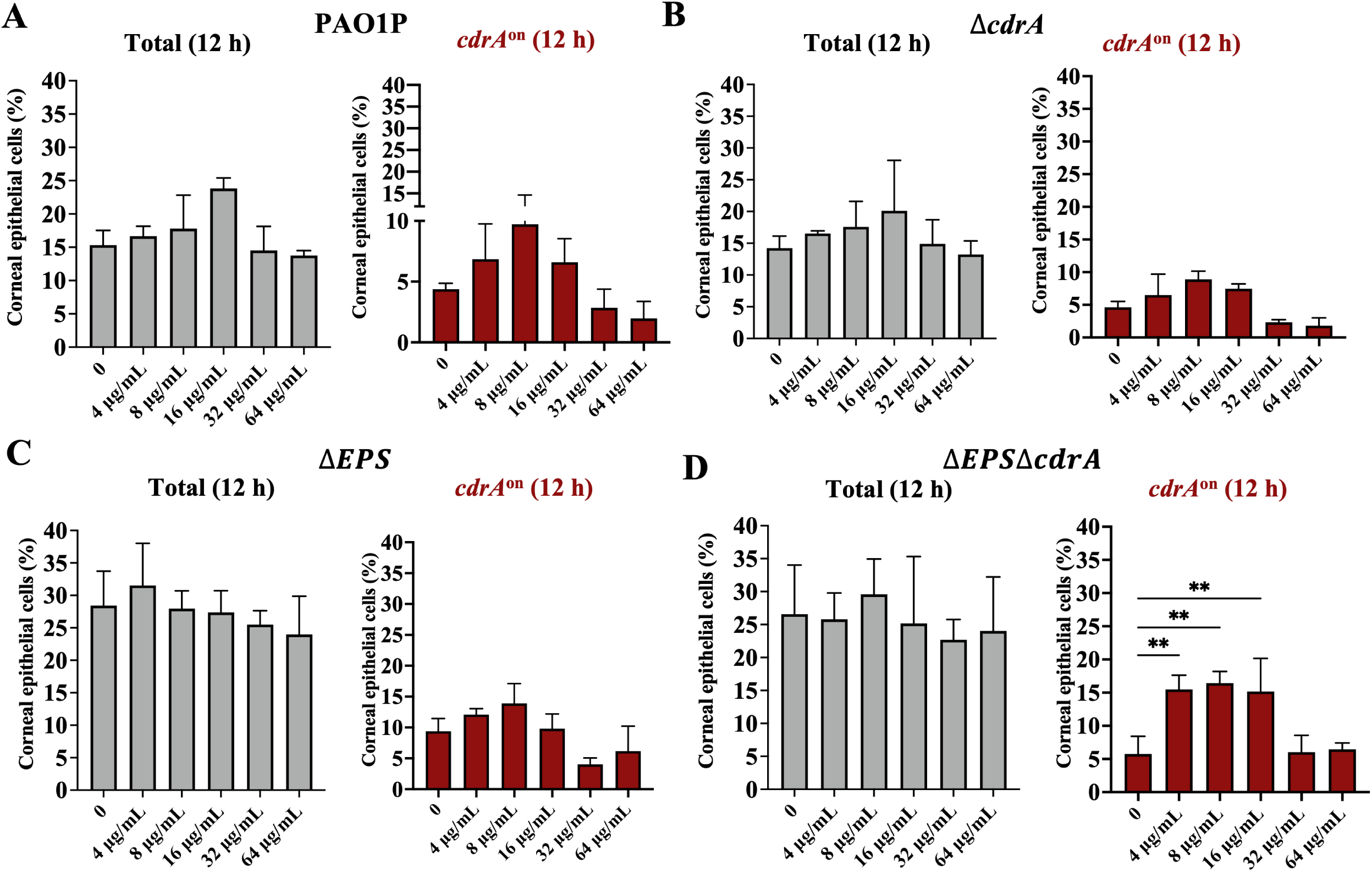
Ofloxacin resistance of vacuolar *P. aeruginosa* does not require *cdrA* or exopolysaccharide. Percentage of human corneal epithelial cells containing intracellular *P. aeruginosa* at 12 h post-infection with (A) PAO1P, (B) PAO1PΔ*cdrA*, (C) PAO1PΔEPS (Δ*pel*Δ*psl*Δ*algD*) or D) PAO1PΔEPSΔ*cdrA*. Total intracellular bacteria were detected using pGFP_arabinose_ (grey bars), and vacuolar *cdrA*^on^ using *cdrA*-GFP (red bars). Extracellular bacteria were killed after 3 h with amikacin (200 μg/mL) and amikacin (200 μg/mL) and ofloxacin added at 3 h (4 – 64 μg/mL = 1 - 16X MIC). Intracellular bacteria persisted up to 16X MIC for wild-type and mutants with a subset of each expressing *cdrA*. * p < 0.05, ** p < 0.01, *** p < 0.001, **** p < 0.0001 (One Way ANOVA with Dunnett’s multiple comparisons test versus untreated control). Data are shown as mean +/- SD of three biological replicates.

As shown in Figures 6 C and D, similar results were obtained for the Δ*EPS* triple mutant lacking all three biofilm associated exopolysaccharides (EPS), and a quadruple mutant additionally lacking CdrA (Δ*EPS*/Δ*cdrA* mutant). In each case the percentage of epithelial cells containing *cdrA*-promotor expressing vacuolar sub-populations at each ofloxacin concentration was similar.

Control experiments confirmed that PAO1F and PAO1P showed similar levels of internalization into human corneal epithelial cells at 4 h post-infection, as did PAO1P relative to its *cdrA* and *EPS* mutants (Supplemental Fig. S2).

### T3SS mutants unable to exit vacuoles show similar ofloxacin resistance to vacuolar located wild-type PAO1

The above data suggested that something about vacuolar localization might promote ofloxacin resistance, independently of expression of classical biofilm-associated factors (e.g. CdrA and EPS) that co-incidentally occur in that location, possibly even a physical feature of the vacuole itself rather than a bacterial-driven mechanism. As a first step toward understanding this, we used a T3SS mutant (PAO1FΔ*exsA*) unable to escape the vacuoles (9–11, 17). Fig. 7 shows time-lapse imaging of human corneal epithelial cells infected with PAO1FΔ*exsA* expressing pGFP_arabinose_ to visualize all intracellular bacteria over time with and without ofloxacin and subsequently quantify percentage of cells containing internalized bacteria. As expected, the Δ*exsA* mutant localized only to vacuoles observed as GFP-expressing puncta within the epithelial cells that increased in fluorescence over time (Fig. 7A). This population survived exposure to ofloxacin at 6 and 12 h post-infection (Fig. 7B, Supplemental Video S3). CLEM was then used to ensure that persistent vacuolar populations were due to live bacteria not residual GFP signal from dead bacteria (42). Results showed the Δ*exsA* mutant localized to perinuclear vacuoles with multiple bacteria in a single membrane-bound compartment (Fig. 7C, inset). Some of these vacuolar bacteria were seen surviving exposure to ofloxacin (1 μg/mL) as shown by bacteria maintaining their cell wall integrity in perinuclear vacuoles (Fig. 7D, inset). Thus, T3SS mutants of *P. aeruginosa* that cannot escape vacuoles can phenocopy the T3SS-off subpopulation of wild-type when infecting corneal epithelial cells with respect to tolerating ofloxacin, supporting the possibility that vacuolar localization itself enables survival.

**Figure 7.**
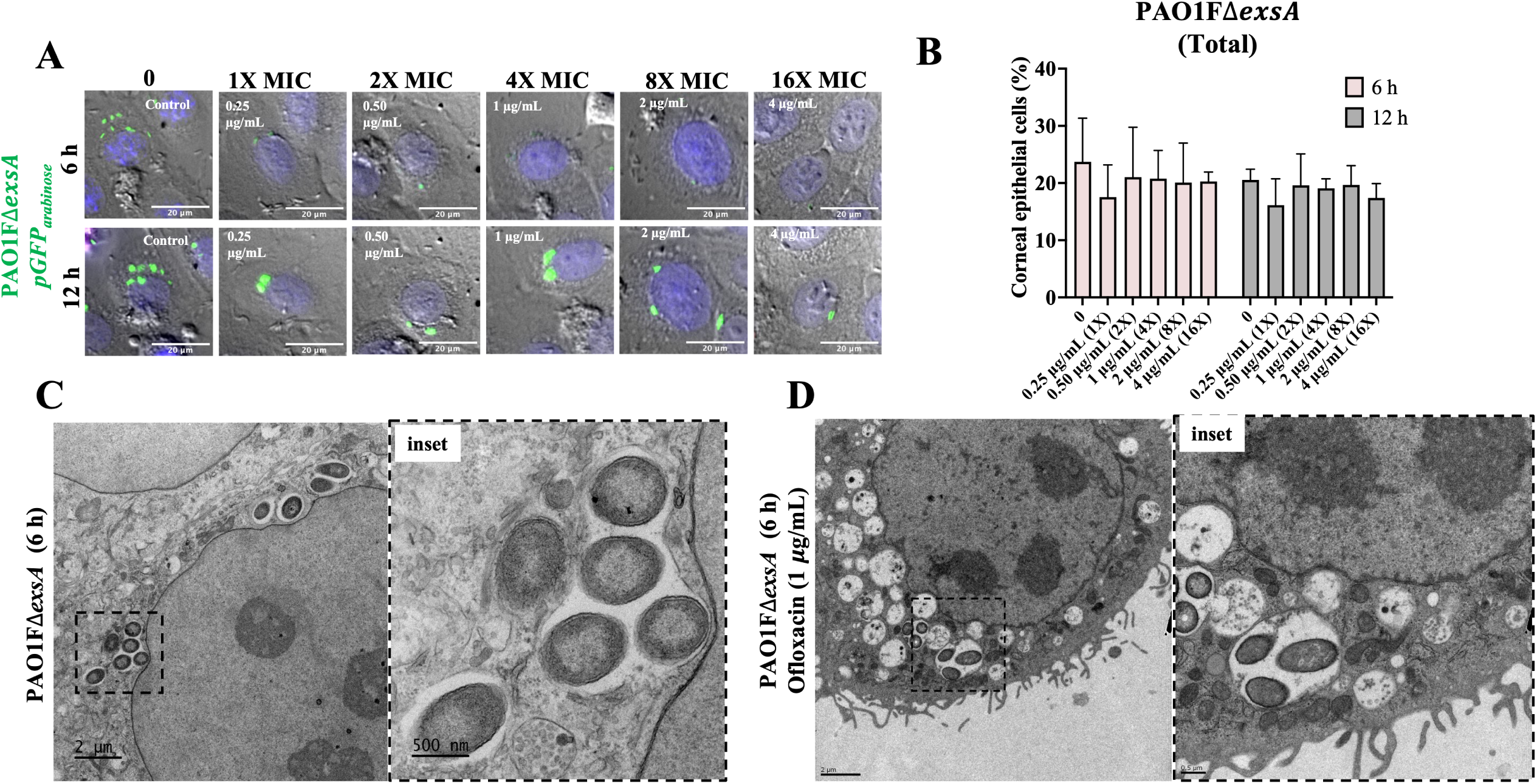
Vacuolar T3SS (Δ*exsA*) mutants are also tolerant to ofloxacin. A) Images of corneal epithelial cells containing PAO1FΔ*exsA* (pGFP_arabinose_) show that all intracellular bacteria are localized to vacuoles with or without ofloxacin (0.25 - 4 μg/mL = 1 - 16X MIC) at 6 and 12 h post-infection. Extracellular bacteria were killed after 3 h with amikacin (200 μg/mL) and then amikacin (200 μg/mL) and ofloxacin added for a further 3 h. B) Percentage of corneal epithelial cells containing at least one bacterial cell at 6 h and 12 h post-infection under the above conditions. Increasing concentrations of ofloxacin failed to eliminate the *exsA* mutant vacuolar population at either time point. There was no significant difference between groups (One-way ANOVA with Dunnett’s multiple comparisons test). C-D) Correlative light electron microscopy to visualize *exsA* mutants in vacuoles in the corneal epithelial cells at 6 h post infection in control (C) and ofloxacin (1 μg/mL) treated (D) cells under the same conditions. Both insets show bacteria restricted to perinuclear vacuoles with evidence of a vacuolar membrane.

### Vacuolar-localized *P. aeruginosa* can drive host cell death

The above experiments showed that vacuolar bacteria can be more tolerant to ofloxacin. To further explore the significance of the vacuolar phenotype, we next examined how it impacts the host cell. Using the same experimental protocols as above, the kinetics of human corneal epithelial cell death during infection were measured using propidium iodide. After cells were infected with PAO1Δ*exsA* (occupies only vacuoles) for 3 h, extracellular bacteria were killed using amikacin (200 μg/mL), then ofloxacin 0 - 4 μg/mL (0 -16X MIC) was added into the media to also kill the cytosolic population. Cells containing intracellular bacteria showed increasing cell death from 4 - 24 h up to maximum of ∼ 40 % of the total population, the kinetics not significantly affected by ofloxacin exposure up to 4 μg/mL (16X MIC) (Fig. 8A). A significant reduction in cell death was only observed when the ofloxacin concentration was increased to 25 μg/mL (100X MIC). While 25 μg/mL (100X MIC) of ofloxacin did not reduce visible GFP-expressing (pGFP_arabinose_) (Fig. 8B), the effect on cell death suggests supraphysiological concentrations of antibiotic are required to inactivate vacuolar-localized populations. This further suggests an active role of intracellular vacuolar bacteria in the outcome of host cell death.

**Figure 8.**
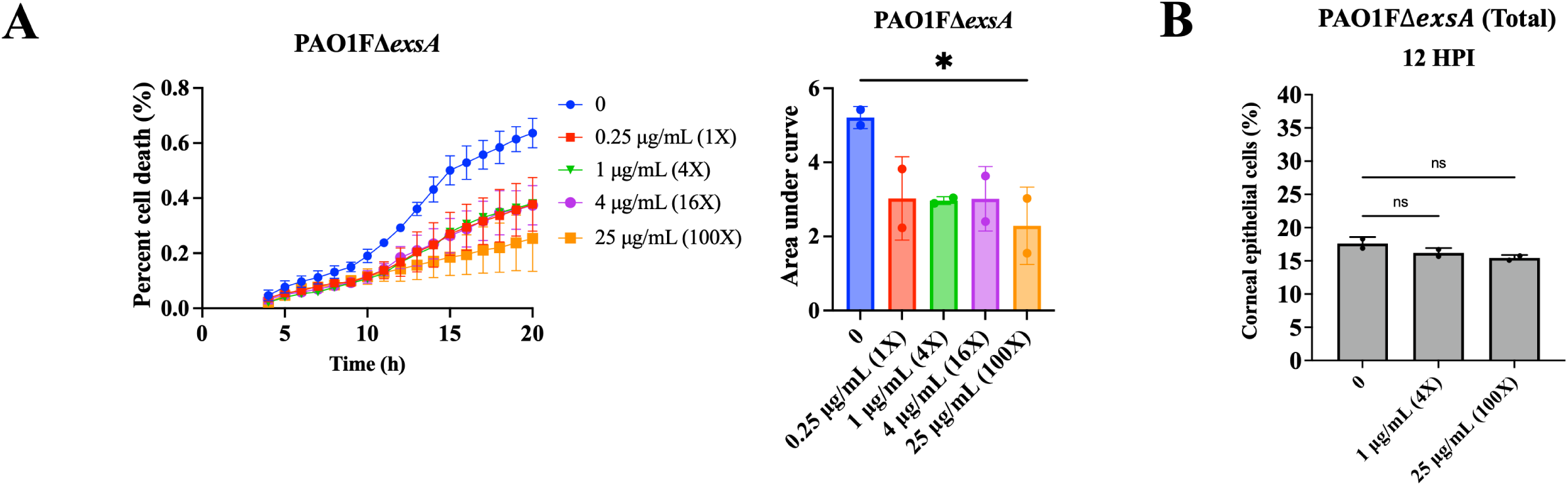
Vacuolar *P. aeruginosa* contribute to host cell death. Quantification of corneal epithelial cell death quantified using the ratio of PI/DAPI, also shown as the area under the curve (AUC) after internalization of *P. aeruginosa* PAO1FΔ*exsA* (A) from 4 to 20 h post-infection. Extracellular bacteria were killed after 3 h with non-cell permeable amikacin (200 μg/mL) and ofloxacin added at 3 h (0.25 - 25 μg/mL = 1 – 100 X MIC) with continued amikacin. Intracellular survival of bacteria shown as percentage of corneal epithelial cells containing a GFP-expressing bacterium (pGFP_arabinose_) treated with 0 (no ofloxacin), 1 μg/mL (4X MIC), 25 μg/mL (100X MIC) at 12 h post-infection (B). The data show that ofloxacin-sensitive and ofloxacin-tolerant vacuolar bacteria both contribute to host cell death. * p < 0.05, ns = Not Significant (One-way ANOVA with Dunnett’s multiple comparisons test versus untreated control).

### Diversification of intracellular phenotypes also occurs during *in vivo* corneal infection

It has been three decades since we initially reported that *P. aeruginosa* can invade corneal epithelial cells *in vivo* in mice, including the first observation that they could be in membrane-bound vacuoles as shown using transmission electron microscopy (TEM) (6). Because TEM required sample fixation, it was unclear if these bacteria were destined to be degraded, to enter the cytoplasm, or to persist in the vacuoles.

More recently, we used intravital confocal/2-photon imaging without fixation to confirm *in vivo* that when *P. aeruginosa* enters the cell cytosol it can colonize the cytoplasm or can traffic to membrane blebs (45). Since the present study revealed intracellular diversification leading to cytosolic and vacuolar bacteria coexisting in the same infected cell population *in vitro*, we next asked if this could also occur *in vivo*. To explore this, we used a murine scarification model that enables *P. aeruginosa* to colonize the epithelium. This was done using Lyz2cre+/mRosa-DTR mice that constitutively express a red fluorescent protein in their cell membranes with along with green/yellow myeloid-derived cells and infecting these mice with mPAO1 constitutively expressing blue-fluorescent protein (pMG055). In this way, we were able to study the localization of infecting bacteria relative to the host cell membranes and differentiate myeloid and non-myeloid derived cell types in the cornea. Live confocal imaging at 15 h post-infection showed three intracellular phenotypes similar in appearance to our *in vitro* observations. These included both cytoplasmic and membrane bleb-contained *P. aeruginosa*, in addition to intracellular aggregates appearing to be vacuoles containing bacteria.

The absence of tissue fixation/sectioning allowed us to additionally collect temporal data in 3D in intact eyes during infection. Doing so, we were able to observe *in vivo* multiple phenomena that we have reported when *P. aeruginosa* infects cultured corneal and other epithelial cells. This included a sub-population in the cytosol demonstrating twitching motility, a second sub-population demonstrating swimming motility inside plasma membrane blebs, and a third population stationary and appearing to be confined to vacuoles (Fig. 9, Supplemental Videos S4, S5, S6). These results suggest that all of the intracellular phenomena that we have reported when *P. aeruginosa* infects cultured cells occur during infection *in vivo*.

**Figure 9.**
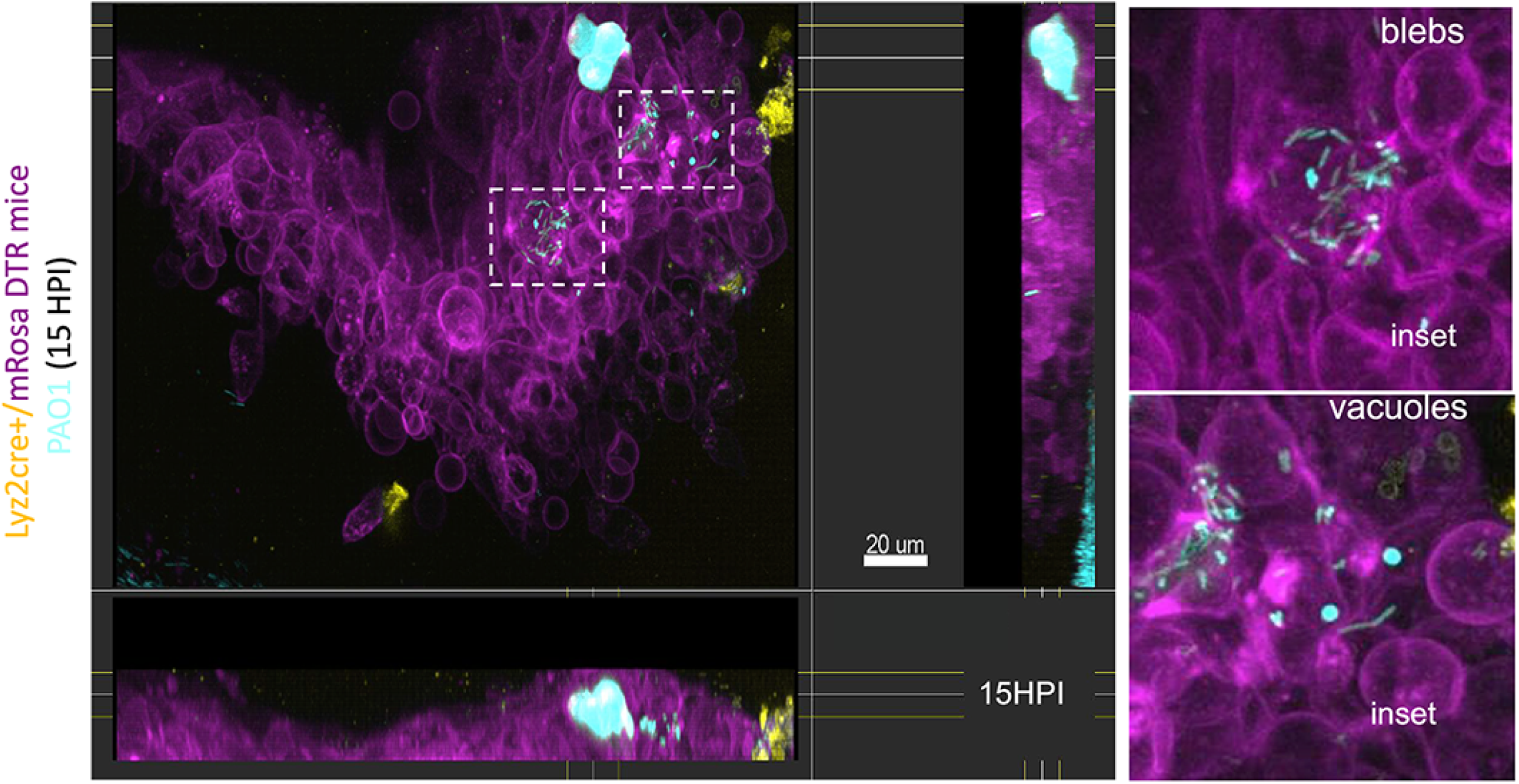
Intracellular cytosolic and vacuolar phenotypes of *P. aeruginosa* both occur *in vivo* during corneal infection. Corneal image of a Lyz2cre+/mRosa-DTR mouse (red fluorescent membranes with green/yellow myeloid-derived cells) at 15 h post-infection with *P. aeruginosa* mPAO1 expressing pMG055 (EBFP2 [blue], pseudo-colored cyan) using the murine scarification model. Confocal images were taken in the middle of the infected cornea scratch on an anesthetized live mouse. Insets show cytosolic bacteria spreading in the cell and/or within membrane blebs, and vacuolar bacteria (blue puncta) within neighboring cells. Side panels show lateral projections of imaging into the epithelium.

## DISCUSSION

Bacteria surviving inside host cells generally orchestrate their localization and replication via secretion of virulence factors to establish a niche in either the cytosol or in subcellular membrane-bound compartments such as vacuoles, secretory vesicles, and endoplasmic reticulum (46–49). Previously, we showed that when *P. aeruginosa* invades epithelial cells it can utilize its type 3 secretion system (T3SS) to escape vacuoles (17). This is followed by rapid replication and dissemination throughout the cytosol, with the T3SS effector ExoS used to avoid autophagy and to construct and occupy host cell plasma membrane blebs (9, 11, 19). We have also reported that *P. aeruginosa* differs from other pathogens by not depending on host cell cytoskeletal elements for intracellular motility, with bacteria instead using their own appendages (25). This includes pilus-driven twitching motility in the cytoplasm and flagella-mediated swimming motility within membrane blebs (11).

The T3SS of *P. aeruginosa* is expressed by only some population members even under inducing conditions (11, 50). Since the T3SS is required for vacuolar escape but not cell entry, T3SS-off subpopulation members remain in vacuoles after being internalized (9–11). Our prior model was that these were destined for degradation in lysosomes, an idea supported by vacuole acidification (10). Here, we instead found that T3SS-off *P. aeruginosa* remained viable over long periods of time in vacuoles, shown using both corneal and bronchial epithelial cells. We also found they could differentiate into a phenotype expressing biofilm-associated factors that are not expressed by their cytoplasmic counterparts. These vacuolar T3SS-off bacteria were often present in cells also containing cytosolic T3SS-on bacteria. Thus, we have identified a novel (third) intracellular niche for *P. aeruginosa* inside epithelial cells in which the bacteria are in a different phenotypic state (T3SS-off, biofilm-producing, non-motile) in addition to being in a different intracellular location (vacuoles). Use of CLEM to simultaneously study the fine structure of the host cells versus location of CdrA-expressing bacteria confirmed that this biofilm protein was expressed by intracellular bacteria residing in vacuoles.

The significance of *P. aeruginosa* residing in epithelial cell vacuoles could potentially have multiple impacts on disease outcome, including presenting challenges to therapeutics. Firstly, we found vacuolar bound bacteria were more tolerant to a cell permeable antibiotic than their cytosolic counterparts. Persistence and antibiotic recalcitrance of vacuole-contained bacteria could be a contributor to why *P. aeruginosa* infections are difficult to treat, and why infection can rebound after antibiotic treatment is ended. Secondly, the data showed the vacuolar population triggered lytic host cell death contingent on their viability, even when the cytosolic population is eliminated using an antibiotic. Indeed, use of an antibiotic at concentrations killing only the cytosolic population actually *promotes* this type of cell death because the cytosolic population uses ExoS to counter it (16). Since lytic cell death releases inflammatory mediators, its inhibition by ExoS likely serves to limit host responses to infection while also preserving the replicative niche. If so, killing cytosolic bacteria with antibiotics that preserve the vacuolar population could favor immune responses/inflammation. How this would play out in an actual infection (e.g. increased bacterial clearance versus damaging inflammation) will require further investigation given the complexities of *in vivo* systems. An additional consideration relates to the design of novel therapies (e.g. targeting bacterial biofilms) if they need to access an intracellular niche enclosed by two host membrane layers.

Intravital confocal imaging of infected mouse corneas was used to explore the location of *P. aeruginosa* within cells after *in vivo* infection. The methods used allowed spatial and temporal resolution, and therefore accurate localization of bacteria with respect to cell membranes over time in addition to their depth within the tissue. Bacteria were detected in the cytosol of infected epithelial cells, some deep within the tissue and others closer to the corneal surface. As we have shown *in vitro*, this included multiple bacteria disseminating through the cell cytoplasm at speeds/pattern aligning with twitching motility, with some cells (also or instead) containing rapidly swimming bacteria inside membrane blebs. A third intracellular bacterial population was detected resembling *P. aeruginosa*-containing vacuoles occurring in cultured epithelial cells. While the fluorescent signal from bacteria packed inside vacuoles merges to prevent resolution of individual bacteria, the circular shape of the emanating fluorescence implies that these bacterial-containing regions are surrounded by a vacuolar membrane.

Mutants lacking the T3SS (Δ*exsA*) remain in vacuoles because they lack the ability to escape the vacuole. These were also found to express biofilm-associated factors, essentially phenocopying T3SS-negative sub-populations of wild-type persisting in vacuoles. While showing that T3SS mutants can be used to study the phenotype, this result also suggests that lack of T3SS expression and subsequently the vacuolar environment may drive biofilm production. Biofilm production in response to the vacuolar environment aligns with our current understanding of biofilms, which can help bacteria tolerate adverse environmental conditions such as nutrient limitation, host immune responses, and exposure to antibiotics (3, 4, 27, 31, 51, 52), and is therefore triggered by such conditions. There are many aspects of the vacuole environment with the potential to trigger the production of biofilm, including vacuolar acidification, expression of intracellular antimicrobial factors such as defensin antimicrobial peptides, reactive oxygen species, and lytic enzymes which all form part of the antimicrobial activity of phagolysosomes (53). Alternatively, biofilm production in vacuoles might relate to the T3SS-off state. *P. aeruginosa* can switch back and forth between T3SS-on/motile/rapid growth state and a T3SS-off/non-motile/biofilm-producing/slow growth/antibiotic-tolerant state, which is controlled by a complex regulatory system (Gac/Rsm pathway) (54). This is often referred to as the acute/chronic switch in infection because the former supports acute infection pathogenesis, and the latter promotes persistence and chronic infection. While features of the cytosolic versus vacuolar niches arising when *P. aeruginosa* diversifies inside cells are reminiscent of the acute and chronic states, the latter are generally believed to occur in different infection types, or at least different time points during infection. Here, we have found the intracellular phenotypes occur simultaneously (including in the same cell) and they both occur relatively early (acute T3SS^on^ 4 h and *cdrA*^on^ 6 h). Related to this, others have reported other situations when the lines between acute and chronic infection phenotypes are blurred (55, 56). More work will be needed to understand the relationship if any between intracellular diversity of *P. aeruginosa* and previously described phenotypic switches.

Our data showed that *P. aeruginosa* in vacuoles expressed biofilm factors known to help it persist in the host and other adverse environments. They were also more antibiotic-tolerant than *P. aeruginosa* found elsewhere in the cell which did not express biofilm factors. Nevertheless, mutants lacking the biofilm protein CdrA and all three biofilm-associated exopolysaccharides retained their ability to persist in vacuoles and tolerate antibiotics, showing that these factors were not required. Accordingly, only a sub-fraction of the vacuolar population expressed *cdrA* when wild-type was used, and the total percentage of cells containing antibiotic-tolerant bacteria exceeded the fraction expressing *cdrA*. While inconsistent *cdrA* expression might relate to different stages of maturation in the vacuole, this result provides further evidence that *cdrA* is not required. Possible explanations for these results include functional redundancy with other biofilm-associated factors, or that different biofilm-associated factors are responsible - possibly even novel biofilm factors produced in this location. Alternatively, persistence/antibiotic tolerance might not be due to biofilm factors. Other possibilities include a reduced growth rate, which can decrease sensitivity to fluoroquinolones (52). Division rates for vacuolar bacteria were obviously slower than the rapid replication occurring in the cytosol (Fig. 3A, D and Fig. 5A), not surprising given space and nutrient limitations, and other adverse conditions expected in vacuoles. Other phenotypic changes that could participate include expression of multidrug efflux pumps or factors that modulate cell wall permeability (57–59), or that vacuolar bacterial sub-population(s) enter a persister state in the face of ofloxacin exposure (60). Rather than being a property of the enclosed bacteria, enhanced tolerance in vacuoles might instead relate to vacuolar properties. While the antibiotic is cell permeable, there will still be some exclusion by the vacuolar membrane. In this regard, the concentration needed to kill cytosolic bacteria (enclosed by the host cell plasma membrane) was somewhat higher than needed to kill extracellular bacteria (∼2-fold). This alone is unlikely to explain the ∼4- to 8-fold additional tolerance of vacuolar bacteria on the other side of a second membrane. However, it is possible that *P. aeruginosa*, like other intracellular vacuolar bacteria, modifies the vacuolar membrane (46, 47, 61, 62). If such modification to the vacuolar membrane limits penetration of antibiotics remains unknown. Since there are many possibilities, pinpointing the mechanisms by which *P. aeruginosa* persists in vacuoles will require further investigation, which could lead to novel strategies for treating antibiotic recalcitrant infections.

While studying the exopolysaccharide (EPS) mutants, we found that they persisted in cells even more efficiently than wild-type, the results showing that they occupied ∼25 – 30 % of cells versus ∼15-20 % of cells for wild-type or *cdrA* mutants able to express EPS (Fig. 6C and D, grey bars). Interestingly, initial internalization (at 4 h) of wild-type and Δ*EPS* mutants was similar (Supplemental Fig. S2) suggesting that greater intracellular persistence of Δ*EPS* mutants at 12 h (Fig. 6C, D) was not due to loss of antiphagocytic activity, the latter being a well-established role for exopolysaccharides in other types of bacteria. However, determining how EPS contributes to modification of bacterial persistence was beyond the scope our aims.

This study further demonstrates the advantage of time-lapse imaging for studying intracellular bacterial populations, particularly for those with a complex lifestyle. For instance, the detection and quantification of fluorescent bacteria using gene-expression reporters allowed the delineation of viable intracellular sub-populations in real-time. Moreover, this methodology is especially of value when studying bacterial persistence after antibiotic treatment as it can detect metabolically-active bacteria that may not be quantifiable using traditional CFU counting, e.g. viable but non-culturable cells. Another advantage of imaging, labeling cellular structures, use of high-resolution methods, and including temporal information, is being able to distinguish intracellular bacteria from antibiotic recalcitrant extracellular bacteria, which can occur if bacteria form extracellular biofilms around the cells being studied (63, 64). A potential *caveat* of our study relates to the arabinose-inducible GFP expression method we used to visualize all intracellular bacteria irrespective of gene expression profile. Recent studies have shown that with some promoters, including the one used here, expression can be impacted by Vfr (65, 66), in which case the number of intracellular bacteria might be greater than those we detected using this method.

In conclusion, this study furthers our understanding of the intracellular lifestyle of *P. aeruginosa* by demonstrating that it diversifies intracellularly to form viable sub-populations in both vacuoles and the cytoplasm of epithelial cells. In these distinct locations they adopt different phenotypes and functions. T3SS-expressing bacteria exit vacuoles, disseminate through the cytoplasm using twitching motility, and use ExoS to replicate rapidly, avoid autophagy, and inhibit cell death to keep the replicative niche alive. Some of these T3SS-expressing cytosolic bacteria also use ExoS to assemble and occupy membrane bleb niches that can subsequently detach, allowing vesicle-contained T3SS-expressing bacteria to travel to distant locations (9). Meanwhile, vacuolar populations in the same or adjacent cells grow slowly, express biofilm-associated factors, tolerate antibiotics, and in the absence of cytosolic cell-mates trigger death of the infected host cell. Since the different phenotypes can occur in the same cell, cooperation is likely and might ensure both short- and long-term interests are met. Other bacteria able to adopt an intracellular lifestyle tend to establish one type of niche in an infected cell type, even if they traffic through various compartments to arrive there. The unique ability of *P. aeruginosa* to diversify intracellularly and assemble multiple niches associated with differential gene expression is likely to contribute to the reasons why it is a highly effective opportunistic pathogen, and why infections caused by it are so difficult to treat.

## MATERIALS AND METHODS

### Cell culture

Human corneal epithelial cells (hTCEpi) (67) were cultured in KGM-2 (Lonza, USA) supplemented with 1.15 mM calcium chloride (high-calcium) to allow differentiation and maintained at 37°C (humidified) in a 5 % CO_2_ incubator. Human bronchial epithelial cells (NuLi-1) were cultured as previously described (19) in BEGM (Lonza, USA) with 1.15 mM calcium chloride and maintained as for hTCEpi. Epithelial cells were grown as monolayers on 24-well tissue culture dishes (MatTek Corporation, Ashland, MA) for wide-field microscopy or plasmid validation respectively, or on 8-well tissue culture dishes (ibidi, Gräfelfing, Germany) or glass-coverslips for use in immunofluorescence.

### Bacterial strains, plasmids, and mutants

*P. aeruginosa* strains and plasmids are shown in Table 1. Bacteria were grown at 37 °C on trypticase soy agar (TSA) (Hardy Diagnostics, Santa Maria, CA) for 16 h before use. For imaging bacteria were transformed with either the T3SS-GFP reporter (pJNE05), c-di-GMP Turbo-RFP reporter (pFy4535)(39), *cdrA*-GFP reporter (pMG078), or pBAD-GFP (arabinose-inducible, pGFP_arabinose_) and grown on media supplemented with gentamicin (100 µg/mL). Bacterial inocula were prepared by resuspending bacterial colonies into sterile PBS to an optical density (O.D) at 550 nm of 1.0 (∼ 2 x 10^8^ CFU/mL).

To construct plasmid pMG078 (*cdrA*-GFP), the pJNE05 empty vector was engineered to have a promoter-less *gfp* and adding MluI and ScaI restriction sites upstream of the *gfp* to create a multiple cloning site for introducing any promoter of interest. Briefly, pJNE05 was prepared as two large PCR products with overlapping ends, with the MluI and ScaI sites added to the primers and excluding the promoter already present in the native pJNE05 vector. Fragment 1 was amplified using the primer pair with engineered restriction sites indicated in bold pJNE05_part1.1a and pJNE05_part1.1b- and Fragment 2 was amplified using pJNE05_part2.1a and pJNE05_part2.1b (Table 2). The two fragments with overlapping ends were assembled by Gibson assembly. The *cdrA* promoter region for the promoter-*gfp* fusion was then amplified using previously published primers (68) with slight modification for our vector backbone (cdrA.1a and cdrA.1b, Table 2). The resulting fragment and promoter-less *gfp*-pJNE05 backbone were digested with MluI and ScaI then fused by T4 DNA ligase to construct the pMG078 vector. The prepared *cdrA*-GFP fusion reporter was confirmed by comparing relative fluorescence intensity in WT (PAO1F) and the PAO1FΔ*fleQ* mutant (11) in the stationary phase of batch culture after ∼12 h growth in TSB at 37°C (normalized to O.D. at 650 nm) (Supplemental Figure S1A). Sensitivity of the *cdrA*-GFP reporter (pMG078) to high levels of c-di-GMP was determined by comparing relative fluorescence intensity in wild-type (PAO1P; low c-di-GMP) and the mutant PAO1PΔ*wspF* (hi-c-di-GMP) over 24 h of growth in TSB at 37°C in a Biotek plate reader at 200 rpm recording every 30 min (Supplemental Figure S1B).

**Table 2.**
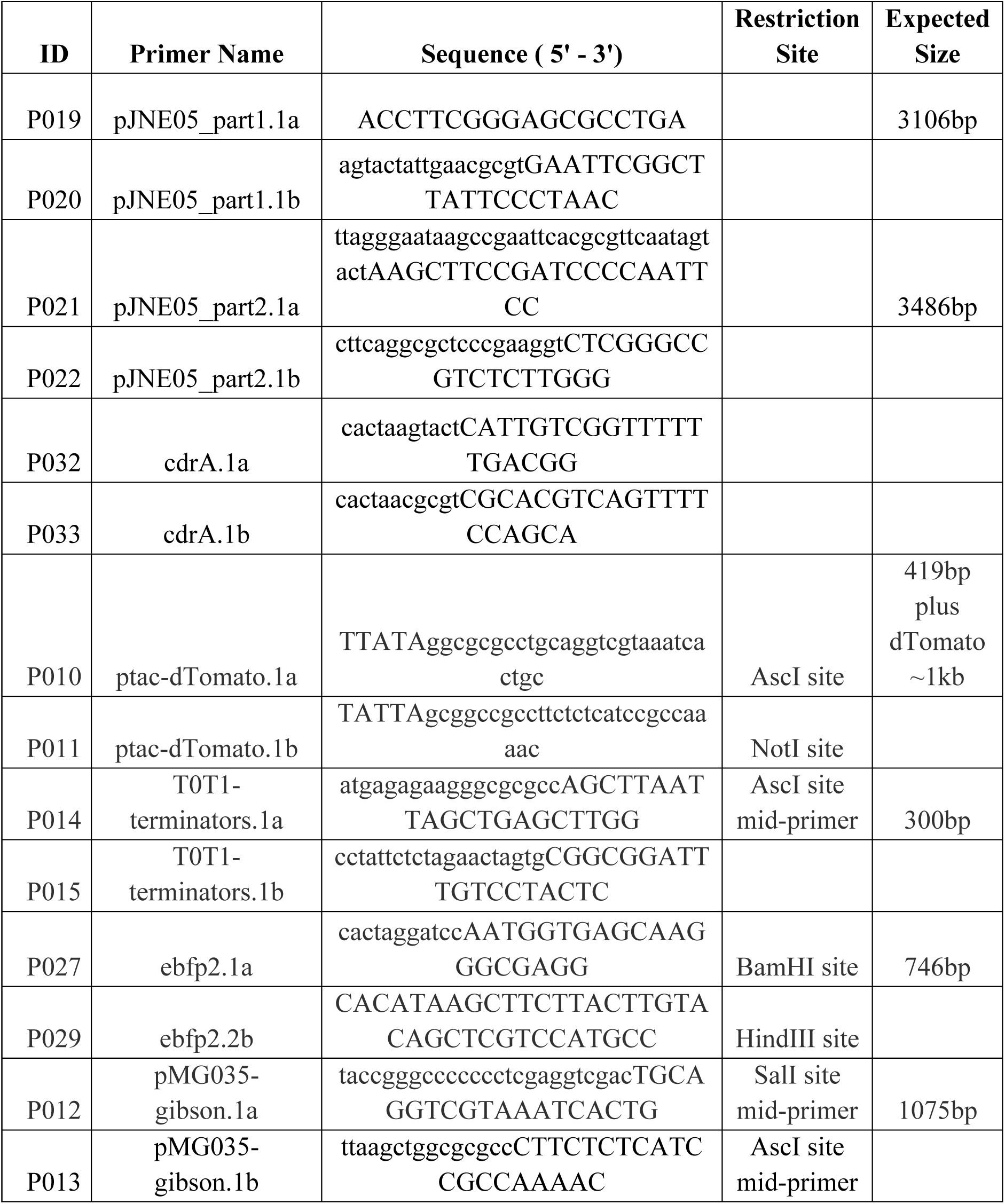
Primers used in this study

To construct pBAD-GFP (pGFP_arabinose_), the promoter region of the pJNE05 was replaced with an arabinose-inducible promoter region. The *exoS* promoter of pJNE05 was excised by digesting with HindIII and EcoRI and replaced with the arabinose-inducible promoter from plasmid pTJ1. Validation of pGFP_arabinose_ *in-vitro* was performed using hTCEpi cultured on 24-well plates. Epithelial cells were inoculated with PAO1F::pGFP_arabinose_ or PAO1FD*exsA*::pGFP_arabinose_ (∼2 x 10^6^ CFU bacteria, MOI of 10) for 3 h. Extracellular bacteria killed by adding amikacin (200 μg/mL) for 30 min. GFP expression was induced after 30 min by adding media containing 1% L-arabinose and the GFP signal recorded hourly for 8 h on a Nikon Ti Eclipse inverted wide-field microscope (Supplemental Fig. S1).

Plasmid pMG055 (constitutive blue fluorescent protein, EBFP2) was constructed as follows. Briefly, to start pMG046 was generated by cloning the tac promoter and dtomato from p67T1 (69) and cloned into the AscI/NotI site of pTJ1. *Ebfp2* from the plasmid pBAD-*ebfp*2 (purchased from Addgene) was then cloned into the BamHI/HindIII site of pMG046 downstream of the tac promoter (replacing dTomato) to create pMG051. Then, the ptac-*ebfp2* construct was amplified from pMG051, plus the T0T1 terminator fragment amplified from pTJ1, and cloned via Gibson assembly into the SalI/BamHI site of promoter-less GFP pJNE05 based plasmid, to yield pMG055 - which has a constitutive EBFP2, and a promoter-less *gfp* gene.

### Bacterial internalization assays

Epithelial cells were inoculated with 5 µl (containing ∼2 x 10^6^ CFU bacteria) of a suspension of *P. aeruginosa* PAO1 wild-type or mutants transformed with fluorescent reporter plasmids (MOI = 10) and incubated for 3 h at 37 °C (internalization period). Epithelial cells were then incubated with cell culture medium (KGM-2 or BEGM + 1.15 mM calcium) supplemented with amikacin (200 μg/mL) for 1 h to kill extracellular bacteria. To explore ofloxacin susceptibility of intracellular bacteria, epithelial cells were exposed to both amikacin (200 μg/mL) and ofloxacin at concentrations ranging from the MIC to 16X MIC. Controls were treated with cell culture media including matched levels of acetic acid diluent. In some experiments, pGFP*_arabinose_* was activated 30 min after extracellular antibiotic was administered using 1 % L-arabinose.

### Microscopy

Live and time-lapse images were captured on a Nikon Ti-E inverted wide-field fluorescence microscope equipped with Lumencor SpectraX illumination source and Okolab Uno-combined controller stage top incubation chamber to maintain heat, humidity, and 5 % CO_2_. Time-lapse images were captured using a CFI Plan Apo Lambda 40X air objective, equipped with differential interference contrast (DIC). Live phase-contrast images were captured using a Plan Apo Lambda Ph3DM 60X oil-immersion objective. For time-lapse, fields were chosen visualizing DIC only to identify areas free of debris; Turbo-RFP and GFP were not observed until time-lapse was completed to avoid bias in field selection. For time-lapse imaging with quantification, eight fields were imaged for each condition. For live phase-contrast without quantification, three fields were imaged for each condition.

### Ofloxacin stock solutions and determination of MIC

Target ofloxacin concentrations from 0.0625 - 8 μg/mL were obtained by dilution of a 10 mg/mL stock solution (prepared in 100 % acetic acid) in cell culture media. Concentrations of 16 - 64 μg/mL were prepared by diluting a 6.4 mg/mL stock solution (prepared in 10 % acetic acid) by 1:400 to 1:100. Similarly, a concentration of 25 μg/mL was prepared by diluting a 2.5mg/mL stock solution (prepared in 10 % acetic acid) by 1:100 dilution. Dilutions were such that acetic acid levels in each assay were below 0.1 %. The MIC of ofloxacin against *P. aeruginosa* wild-type and mutant strains was determined by inoculating ∼1.5 x 10^8^ CFU of bacteria in tryptic soy broth into a 96-well plate at 37 °C, then incubating overnight (16 h) in the presence of ofloxacin at the above concentrations. The MIC was determined as the lowest concentration that inhibited bacterial growth measured by absorbance at 550 nm.

### Immunofluorescence

Epithelial cells were cultured and inoculated with bacteria as described above. After 3 h, KGM-2 (+ 1.15 mM calcium) containing amikacin (200 μg/mL) was added to kill extracellular bacteria for 3 h at 37 °C or amikacin with ofloxacin (up to 16X MIC) added to also challenge intracellular bacteria. At 6 h post-inoculation, cells were washed with PBS, and fixed in fresh 4 % paraformaldehyde (PFA) (Sigma-Aldrich) for 10 min. After an additional PBS wash, cells were permeabilized for 10 min using 4 % PFA containing 0.1 % Triton X-100. Cells were quenched with aldehydes in 150 mM glycine (Sigma-Aldrich) for 10 min, washed in PBS, and blocked for 1 h at room temperature in 0.7 % fish skin gelatin (Sigma-Aldrich). All *P. aeruginosa* bacteria were labeled with anti-*Pseudomonas* antibody (Abcam, #ab74980, primary) with Alexa-Fluor 555 (Sigma, secondary) for 1 h. For all preparations: cells were washed with PBS, ProLong™ Diamond Antifade Mounting Medium (ThermoFisher Scientific) was applied and samples were sealed with a coverslip and imaged. Epithelial cell nuclei were stained with NucBlue® Live ReadyProbes® Reagent (ThermoFisher Scientific).

### Correlative Light and Electron Microscopy (CLEM)

The hTCEpi were grown to 70 % confluence on 35 mm grided Mattek™ dishes with a glass bottom by adding ∼ 6 x 10^5^ cells in KGM-2 media containing 1.15 mM calcium chloride and incubating overnight in a cell culture incubator. Prior to infection, Hoechst dye was added at a concentration of 3 μl/mL to stain the nuclei. Epithelial cells were infected with bacteria at an MOI of 10 and allowed to progress for 3 h. At this time, 1:1 dilution of prewarmed amikacin (400 μg/mL) in KGM-2 + 1.15 mM calcium was added for 30 min followed by ofloxacin to a final concentration of 1 μg/mL for 3 h (total infection time = 6 h). At 6 h, 2 mL of media was removed leaving 1 mL behind to prevent cells drying. Cells were washed three times with the sequential addition of 1 mL of cell culture media always leaving behind 1 mL. Cells were fixed with 1 mL of fixative media in KGM-2 (8 % electron microscopy grade paraformaldehyde [PFA]) for a minimum of 30 min. All fluorescent images were obtained in PFA-containing media. Following fluorescence imaging, cells were fixed in 2.5% glutaraldehyde and 2.5% paraformaldehyde in 0.1M sodium cacodylate buffer, pH 7.4 (EMS, Hatfield, PA, USA). Samples were rinsed 3 times (5 min each) at room temperature) in 0.1M sodium cacodylate buffer, pH 7.2, and immersed in 1% osmium tetroxide with 1.6% potassium ferricyanide in 0.1M sodium cacodylate buffer for 30 min. Samples were rinsed 3 times (5 min each) at room temperature in buffer and briefly washed once with distilled water (1 min) at room temperature. Samples were then subjected to an ascending ethanol gradient followed by pure ethanol. Samples were then progressively infiltrated (using ethanol as solvent) with Epon resin (EMS, Hatfield, PA, USA) and polymerized at 60 °C for 24 - 48 h. Care was taken to ensure only a thin amount of resin remained within the glass bottom dishes to enable the best possible chance for separation of the glass coverslip. Following polymerization, the glass coverslips were removed using ultra-thin Personna razor blades (EMS, Hatfield, PA, USA) and liquid nitrogen exposure, as needed. Correlative Light and Electron Microscopy (CLEM), was performed to visualize specific cells of interest. Regions of interest, identified by the gridded alpha-numerical labeling on the plates were carefully removed, precisely trimmed to the cell of interest, and mounted on a blank resin block with cyanoacrylate glue for sectioning. Serial thin sections (80 nm) were cut using a Leica UC6 ultramicrotome (Leica, Wetzlar, Germany) from the surface of the block until approximately 4-5 microns within to ensure complete capture of the cell volumes. Section-ribbons were then collected sequentially onto formvar-coated slot- or 50-mesh grids. The grids were post-stained with 2% uranyl acetate followed by Reynold’s lead citrate, for 5 min each. Sections were imaged using a FEI Tecnai 12 120kV TEM (FEI, Hillsboro, OR USA) and data recorded using either a Gatan US1000 CCD with Digital Micrograph 3 or a Gatan Rio 16 CMOS with Gatan Microscopy Suite software (Gatan Inc., Pleasanton, CA, USA).

### Murine infection model

All procedures involving animals were carried out in accordance with the standards established by the Association for the Research in Vision and Ophthalmology, under a protocol AUP-2019-06-12322 approved by the Animal Care and Use Committee, University of California Berkeley. This protocol adheres to PHS policy on the humane care and use of laboratory animals, and the guide for the care and use of laboratory animals. For *in-vivo* imaging of corneal infection by *P. aeruginosa*, 8-10 week old female C57BL/6 Lyz2cre+/mRosa-DTR mice (F1 cross) were used since they express red fluorescent membranes and green/yellow myeloid-derived cells. The cornea scarification model was used to establish infection as previously described (70, 71). Briefly, mice were anesthetized for 4 h by ketamine-dexmedetomidine injection (ketamine 80-100mg/Kg and dexmedetomidine 0.25-0.5 mg/Kg) and the cornea of one eye scratched in parallel three times with a sterile 26 G needle, then inoculated with 5 μL of wild-type mPAO1-EBF2 (pMG055; constitutive blue fluorescent protein) in suspension containing ∼ 10^9^ CFU of bacteria at 1 h intervals for 4 h while under continued anesthesia ketamine-dexmedetomidine injection. After 4 h, mice were woken up with anesthesia reversal agent Atipamezole via injection (2.5 – 5 mg/Kg) and remained under supervision for 10 h. At 15 h, mice were re-anesthetized ketamine-dexmedetomidine injection (ketamine 80-100mg/Kg and dexmedetomidine 0.25-0.5 mg/Kg) and infected corneas imaged in the live animals by confocal microscopy.

### Statistical analysis

Prism software was used for numerical data analysis. Data were expressed as a mean +/- standard deviation (SD). Two group comparisons were performed using Student’s t-Test, and multiple group analysis was performed using One-way ANOVA with Dunnett’s multiple comparisons test. *P* values less than 0.05 were considered significant.

## Acknowledgements

Thanks to Dr. Alain Filloux (Imperial College London, UK), Dr. Fitnat Yildiz (University of California Santa Cruz, CA) and Dr. Matthew Parsek (University of Washington, WA) for providing *P. aeruginosa* wild-type strains, respective mutants and plasmid constructs. Thanks to Dr. Danielle Robertson (University of Texas Southwestern, TX) for providing the telomerase-immortalized human corneal epithelial cells. Thanks to Dr. Danielle Jorgens and Ms. Reena Zalpuri at the University of California Berkeley Electron Microscope Laboratory for expert advice and assistance with electron microscopy.

## Contributions

NGK, VN, ARK, MG, MM, TY, DE and SF designed experiments; NGK, VN, ARK, EJ, MH, MG and MM performed the experiments; NGK, VN, ARK, MG, MM, TY, DE and SF analyzed and interpreted the data; NGK, VN, ARK, DE and SF wrote the manuscript; DE and SF supervised the study.

## Supplemental Information

**Supplemental Figure S1.** Validation of *cdrA*-GFP and *pGFP_arabinose_* reporters. A) Maximum fluorescence intensity (Relative Fluorescence Units, RFU) of the *cdrA*-GFP reporter (pMG078) in PAO1F and PAO1FΔ*fleQ* normalized to growth (O.D. at 650 nm) in the stationary phase (after ∼12 h growth in TSB). B) Responsiveness of pMG078 to c-di-GMP levels. GFP fluorescence intensity normalized to O.D. at 550 nm of PAO1P (low-c-di-GMP) (black circles) and PAO1PΔ*wspF* (hi-c-di-GMP) (green squares) over 24 h growth at 30 min intervals. C) Inducible-GFP expression by intracellular PAO1F or its Δ*exsA* mutant (vacuolar) in human corneal epithelial cells using 1 % arabinose. Extracellular bacteria were killed with amikacin 200 μg/mL at 3 h post-infection and cells imaged at 1 h intervals from 4 - 7 h. Color gradient shows the intensity of GFP expression. D) Upper panels show growth (O.D. at 550 nm) of wild-type *P. aeruginosa* parent strains transformed with *pGFP*_arabinose_ over 16 h. PAO1P (blue) or PAO1F (red) alone (closed symbols) or with 1 % arabinose (open symbols). Lower panels show concomitant recording of GFP intensity.

**Supplemental Fig. S2.** Bacterial internalization control experiments. Invasion of wild-type PAO1F and PAO1P into human corneal epithelial cells at 4 h post-infection (left panel) compared to PAO1P (WT) and its mutants in *cdrA*, EPS, or both (right panel) as determined by amikacin exclusion assay. Bars represent mean +/- SD of three biological replicates. ns = Not Significant (Student’s t-Test, left panel; One-way ANOVA with Dunnett’s multiple comparisons test, right panel).

## Supplemental Video Legends

**Supplemental Video S1**. Time-lapse imaging of human corneal epithelial cells infected with PAO1F expressing pJNE05 (T3SS-GFP) showing T3SS^on^ bacteria from 4 - 24 h post-infection. Cells were infected with bacteria at an MOI of 10 and incubated for 3 h. At 3 h, antibiotic-containing media was added to kill extracellular bacteria only (amikacin, 200 μg/mL) or extracellular and intracellular bacteria (amikacin, 200 μg/mL and ofloxacin, 0.25 - 4 μg/mL [1-16X MIC]). Cells were imaged at 1 h intervals for 20 h. Blue = Hoechst staining of nuclei (added during the incubation period), Red = Propidium Iodide staining (added at 3 h to stain dead host cells), Green = bacteria expressing GFP.

**Supplemental Video S2.** Time-lapse imaging of human corneal epithelial cells infected with PAO1F expressing pMG078 (*cdrA*-GFP) showing *cdrA*^on^ bacteria from 4 - 24 h post-infection. Cells were infected with bacteria at a MOI of 10 and incubated for 3 h. At 3 h antibiotic containing media were added to kill extracellular bacteria only (amikacin, 200 μg/mL) or extracellular and intracellular bacteria (amikacin, 200 μg/mL and ofloxacin, 0.25 - 4 μg/mL [1-16X MIC]). Cells were imaged at 1 h intervals for 20 h. Blue = Hoechst staining of nuclei (added during incubation period), Red = Propidium Iodide staining (added at 3 h to stain dead host cells), Green = bacteria expressing GFP.

**Supplemental Video S3.** Time-lapse imaging of human corneal epithelial cells infected with PAO1FΔ*exsA* expressing pGFP_arabinose_ showing total intracellular vacuolar bacteria from 4 - 24 h post-infection. Cells were infected with bacteria at a MOI 10 and incubated for 3 h. At 3 h, antibiotic containing media was added to kill extracellular bacteria only (amikacin, 200 μg/mL) or extracellular and intracellular bacteria (amikacin, 200 μg/mL and ofloxacin, 0.25 - 4 μg/mL [1-16X MIC]). Cells were imaged at 1 h intervals for 20 h. Blue = Hoechst staining of nuclei (added during incubation period), Red = Propidium Iodide staining (added at 3 h to stain dead host cells), Green = bacteria expressing GFP.

**Supplemental Video S4.** Live confocal imaging of the cornea of a Lyz2cre+/mRosa-DTR mouse infected for 15 h with mPAO1-pMG055 (EBFP2 [blue], pseudo-colored cyan) using the corneal scarification model. Video shows intracellular cytosolic bacteria that have occupied membrane blebs and distinct blue puncta suggesting bacteria located in vacuoles. Horizontal projection of the X-Y plane to the right and Y-Z plane in the lower panel.

**Supplemental Video S5.** Live confocal imaging of the cornea of a Lyz2cre+/mRosa-DTR mouse infected for 15 h with mPAO1-pMG055 (EBFP2 [blue], pseudo-colored cyan) using the corneal scarification model. Video shows an inset of cytosolic bacteria that have occupied membrane blebs some demonstrating rapid swimming motility.

**Supplemental Video S6.** Live confocal imaging of the cornea of a Lyz2cre+/mRosa-DTR mouse infected for 15 h with mPAO1-pMG055 (EBFP2 [blue], pseudo-colored cyan) using the corneal scarification model. Video shows an inset of blue puncta indicative of bacteria within vacuoles.

**Animal Statement**. All procedures involving animals were carried out in accordance with the standards established by the Association for the Research in Vision and Ophthalmology, under a protocol AUP-2019-06-12322 approved by the Animal Care and Use Committee, University of California Berkeley, an AAALAC accredited institution. This protocol adheres to PHS policy on the humane care and use of laboratory animals, and the guide for the care and use of laboratory animals.

